# Lamin A redistribution mediated by nuclear deformation determines dynamic localization of YAP

**DOI:** 10.1101/2020.03.19.998708

**Authors:** Newsha Koushki, Ajinkya Ghagre, Luv Kishore Srivastava, Chris Sitaras, Haruka Yoshie, Clayton Molter, Allen J. Ehrlicher

**Affiliations:** Department of Bioengineering, McGill University, Montreal H3A 0E9; Department of Anatomy and Cell Biology, McGill University, Montreal H3A 0C7

**Keywords:** YAP, nucleus, contractile work, TFM, substrate stiffness, cytoskeleton, LINC, deformation, mechanotransduction, mechanosensation

## Abstract

YAP is a key mechanotransduction protein with essential roles in diverse physiological processes. Dysregulation in YAP activity is associated with multiple diseases such as atherosclerosis, fibrosis, and cancer progression. Here we examine the physical stimuli that regulate dynamic YAP translocation to the nucleus. Through a combination of biophysical studies, we demonstrate that YAP localization is insensitive to cell substrate stiffness, but strongly determined by cellular contractile work, which in turn deforms the nucleus. We show that nuclear deformation from LINC-mediated cytoskeletal contractility or extracellular osmotic forces triggers YAP nuclear localization. By modulating the expression of lamin A and nuclear stiffness, we illustrate that nuclear rigidity modulates deformation-mediated YAP nuclear localization. Finally, we show that nuclear deformation causes relocalization of lamin A from the nuclear membrane to the nucleoplasm, and this is essential in allowing YAP to enter the nucleus. These results reveal key physical nuclear deformation mechanics that drive YAP nuclear import.

## Introduction

Sensing and correctly responding to mechanical signals are essential aspects of biology. There are diverse mechanisms enabling cells to sense various mechanical stimuli, including ECM rigidity (Aragona et al., 2013; Cui et al., 2015; Discher et al., 2005; Elosegui-Artola et al., 2016; Engler et al., 2006; Ghibaudo et al., 2008; Kohn et al., 2015), dynamic stretching (Cui et al., 2015), cytoskeletal strain (Benham-Pyle et al., 2015; Ehrlicher et al., 2011) and compression (Guo et al., 2017). Many mechanosensory mechanisms regulate transcription factors, which in turn dictate fundamental aspects of cellular function, homeostasis, and tumorigenesis (Cho et al., 2017b; Driscoll et al., 2015; Dupont et al., 2011; Humphrey et al., 2014). Yes Associated Protein (YAP) is a crucial transcription factor that mediates the interplay between cellular mechanics and signaling cascades underlying gene expression, cell proliferation, differentiation fate decisions, and organ development (Dupont et al., 2011; Kofler et al., 2018; Pavel et al., 2018; Piccolo et al., 2014; Varelas, 2014; Zhao et al., 2010; Zhao et al., 2007). Thus, the spatio-temporal localization of YAP provides critical information about the regulatory state of the cell. Increased YAP activity can cause abnormal and uncontrollable cell proliferation and invasiveness, leading to multiple diseases, including fibrosis and diverse cancers as a result of activating genes associated with oncogenic transcription factors (Donato et al., 2018; Lee et al., 2019; Varelas, 2014; Zanconato et al., 2016b; Zhao et al., 2010).

Microenvironment mechanics appears to influence YAP localization, with several studies reporting that YAP nuclear localization and activity tends to increase with increasing substrate stiffness (Das et al., 2016; Dupont et al., 2011; Elosegui-Artola et al., 2016; Fischer et al., 2016; Nardone et al., 2017). This has led to the hypothesis that substrate stiffness directly influences YAP localization and its resulting effects such as proliferation and differentiation (Dupont et al., 2011; Huebsch et al., 2010; Oliver-De La Cruz et al., 2019; Rammensee et al., 2017). However, the abrogation of these effects by cytoskeletal disruption demonstrates that YAP activity is more directly related to cytoskeletal processes and contractility rather than to substrate mechanics (Das et al., 2016; Dupont et al., 2011; Fischer et al., 2016; Shiu et al., 2018). Elosegui-Artola et al demonstrated that disrupting the actin-LINC complex (Linker of Nucleoskeleton and Cytoskeleton) attenuates YAP activity’s correlation with substrate stiffness, further suggesting a role of cytoskeletal contractility in YAP activity (Elosegui-Artola et al., 2017). However, how contractile forces vary during cell movement, and how they actually determine dynamic movement of YAP in real-time remains unclear, but they are hypothesized to be related to nuclear mechanosensing. Recent work which demonstrates that direct application of force to the nucleus is sufficient to regulate YAP activity reinforces the idea that nuclear deformation is a key mechanism of YAP regulation (Elosegui-Artola et al., 2017). These findings suggest that nuclear deformability and nuclear deformation have essential roles in cells correctly responding to cues from their mechanical environment.

Among the proteins underlying the inner nuclear membrane, lamin A is known as one of the crucial intermediate filament proteins that confers physical support of the nucleus and modulates nuclear stiffness (Athirasala et al., 2017; De Vos et al., 2011; Lammerding, 2011; Lammerding et al., 2006; Lammerding et al., 2004; Olins et al., 2008; Pajerowski et al., 2007). Mutations in lamin A are associated with impaired nuclear mechanotransduction resulting in a variety of human diseases (Bonne et al., 1999b; De Sandre-Giovannoli et al., 2003; Eriksson et al., 2003; Fatkin et al., 1999; Lammerding et al., 2004). Lamin A expression level also modulates nuclear rigidity; suppression of lamin A softens the nuclei and increases nuclear deformability (Guilluy et al., 2014; Hanson et al., 2015; Lammerding et al., 2004), whereas lamin A overexpression increases nucleus’ stiffness (Hanson et al., 2015; Harada et al., 2014). Lamin A expression levels can be affected by cell division (Moir et al., 2000) and stem cell differentiation (Schirmer and Gerace, 2004). Previous studies have also illustrated an interplay between lamin A expression and substrate stiffness (Buxboim et al., 2017; Buxboim et al., 2010; Swift and Discher, 2014; Swift et al., 2013). Soft substrates where the cells have limited spread area, promote lamin A turnover, phosphorylation and subsequent lamin A degradation. Whereas stiff substrates stabilize lamin A as a result of larger forces being transduced to the nucleus, leading to nuclear tension and conformational change in lamins which prohibits access of kinases (Athirasala et al., 2017; Bertacchini et al., 2013; Cho et al., 2017a; Goldman et al., 2002; Miroshnikova et al., 2017; Swift et al., 2013). The similar impacts of mechanical cues on lamin A expression levels and YAP activation, and the correlation of these proteins with stem cell differentiation and cell division suggest a relationship between lamin A, nuclear mechanics, and YAP mechanosensing, however, no such link has been established.

## Results

### YAP nuclear localization is dynamic and independent of cell spread area and substrate stiffness

To examine the dynamics of YAP localization, we transfected living NIH 3T3 cells with EGFP-YAP (pEGFP-C3-hYAP1, Addgene, plasmid #17843) and with EBFP2-Nucleus-7 (nuclear localization signal, Addgene, plasmid #55249) to visualize the nucleus. Similar to previous studies, our principal metric for YAP activity is the ratio of fluorescence of EGFP-YAP in the nucleus to EGFP-YAP in the cytoplasm and is henceforth referred to as the YAP Ratio (YR) (Figures 1A-1C, Figure S1A). To examine how cellular interactions with the substrate determine endogenous YAP and EGFP-YAP localization, we quantified the YR as cells spread on polydimethylsiloxane (PDMS) substrates with different Young’s moduli (0.3 and 48 kPa) (Au - Yoshie et al., 2019; Yoshie et al., 2018). Both transfected and immunostained cells displayed a broad range of YRs on soft PDMS, stiff PDMS, and glass substrates without a stiffness trend (Figures 1A-1C, Figure S1A). We observed both nuclear (YR>1.5) and cytoplasmic (YR<1) localization of endogenous YAP and EGFP-YAP independent of substrate rigidity.

**Figure 1.**
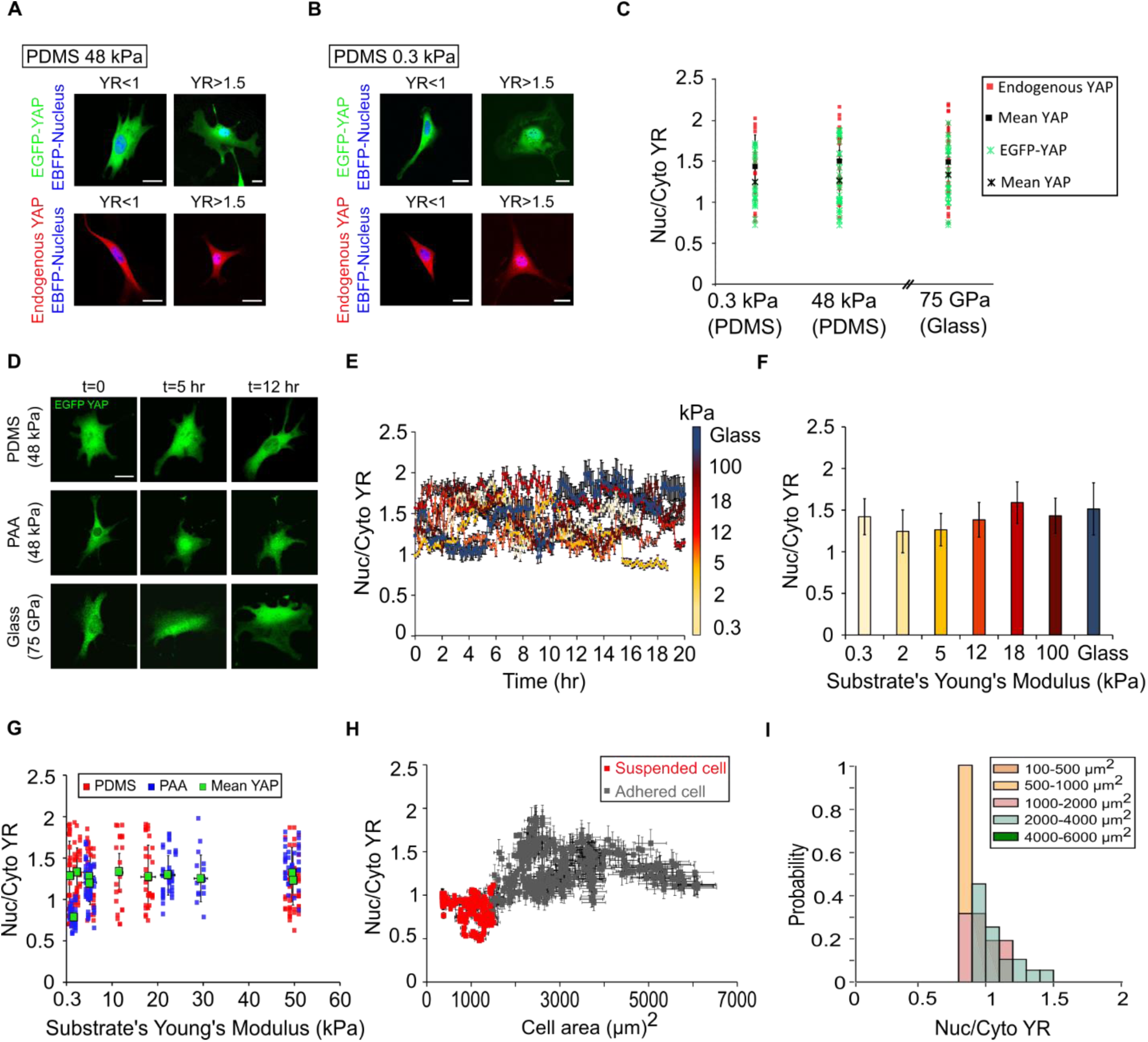
YAP localization is dynamic overtime and independent of substrate stiffness or cell spread area. A) Example image of EGFP-YAP and endogenous YAP merged with EBFP-NLS nucleus on stiff PDMS and, B) soft PDMS, C) YR variation on soft, stiff PDMS, and glass for EGFP and endogenous YAP. Black squares and black stars are mean YR values for EGFP-YAP and endogenous YAP, respectively on each substrate (n>15 cells per each condition), D) Example of EGFP-YAP variation during cell movement on PDMS, PAA, and glass over time, E) Quantification of YR during cells movement on PDMS substrates with different stiffnesses and on the fibronectin-coated glass, F) Time average of YR in the same condition as in (E), G) YR variation for NIH 3T3 cells seeded on different PDMS (shades of red) and PAA (shades of blue) substrates with different stiffnesses (n>20 cells per each condition), H) Quantification of YR and cell spreading area before (red) and during cells attachment (gray) on 5 kPa PDMS substrate (n>10 cells), I) YAP distribution based on cell spread area for an example cell before and during adhesion to PDMS with the same modulus. Scales bars are 20 µm. Error bars indicate standard deviation (SD).

We also monitored the EGFP-YAP distribution during cell movement on PDMS substrates finding that EGFP-YAP localization is highly dynamic in time, with no stiffness correlation over time across diverse PDMS substrate moduli (0.3, 2, 5, 12, 18, 100 kPa) and fibronectin-coated glass (Figures 1D-1F, Figures S1B and S1C). The time-averaged YR on stiff substrates appeared identical to compliant substrates, suggesting that neither the magnitude of YAP localization nor the shuttling frequency is set by substrate stiffness (Figures 1D-1F).

Previous cell substrate studies on polyacrylamide (PAA) have reported a positive correlation between the YR and substrate stiffnesses; YAP cytoplasmic localization was only observed in round cells with small spread area on soft PAA substrates with moduli less than 1 kPa (Das et al., 2016; Dupont et al., 2011; Elosegui-Artola et al., 2017), whereas 5 kPa was identified as a critical modulus for high YR (Aragona et al., 2013; Dupont et al., 2011; Elosegui-Artola et al., 2017). Using PDMS substrates, we observed that the YR did not correlate with substrate stiffness (Figures 1F and 1G). One possible explanation may be ascribed to non-mechanical variations in PAA substrates absent on PDMS (Trappmann et al., 2012).

To examine the impact of PDMS or PAA on YAP localization, we prepared PAA gels with uniform polymer mass and tuned the substrates stiffness by varying the crosslinker concentration. Similar to previous studies, cells on very compliant PAA gel with a Young’s modulus of 1 kPa were round with low spread area and low YAP activity (YR<1) (Figure 1G, Figures S1D, S1E and S1I). However, cells on very compliant PDMS with a Young’s modulus of 300 Pa displayed a wide range of spread areas and random distributions of YAP between their nuclei and cytoplasms, similar to those on the stiff PDMS (Figures 1E-1G, Figures S1F-S1H). Cells on stiff PAA substrates with moduli of 5, 20, and 50 kPa displayed highly dynamic EGFP-YAP localization with diverse spread areas similar to those we observed on PDMS substrates (Figures 1D and 1G, Figures S1D, S1E, S1J and S1K). These results suggest that YAP nuclear localization may be reduced on very soft (E≤1kPa) PAA substrates due to a lack of cell spreading.

To examine the roles of substrate adhesion and cell spread area on YAP translocation, we also tracked YAP localization in round suspended cells (Figure 1H). Here, during initial cell attachment, we consistently measured low YRs in round cells with small spread area (Figures 1H and 1I). As soon as those suspended cells attached and spread on the 5 kPa PDMS substrate, YAP distributed randomly between the nucleus and cytoplasm with no clear correlation with cell spreading (Figures 1H and 1I). Our results suggest that only in the case of rounded exceptionally small cell areas (<∼1000 µm^2^) is YAP consistently in the cytoplasm.

The apparent random spatio-temporal localization of YAP on both PAA and PDMS substrates, coupled with an absence of correlation between YAP localization and substrate stiffness or cell spreading implies that another mechanism may impact YAP localization. Actin stress fibers and intracellular forces may more directly regulate YAP localization (Bouzid et al., 2019; Cho et al., 2017a; Elosegui-Artola et al., 2017; Guilluy and Burridge, 2015; Martino et al., 2018; Shiu et al., 2018). We thus suspected that dynamic changes in contractile stress may lead to dynamic and correlated changes in YAP localization.

### YAP nuclear localization increases with cell contractility

We employed Traction Force Microscopy (TFM) (Butler et al., 2002) to track single-cell contractility in time on PDMS substrates with specified moduli, and compared these metrics with the dynamic YR; we found that as the contractile state of the cell changed in time, it was temporally correlated with the YR (Figures 2A-2E, Figures S2A-S2H). We then compared the instantaneous contractile states of single cells with their YRs over a broad range of substrate stiffnesses, finding that both traction stress and substrate bead displacement correlated with YRs of the corresponding cell (Figures 2F and 2G). These data separated into different correlated populations as a function of substrate stiffness. When we examined YR as a function of cell contractile work for different PDMS substrates, we found all data collapsed onto a single curve (Figure 2H), illustrating that cell contractile work appears to relate to YAP localization.

**Figure 2.**
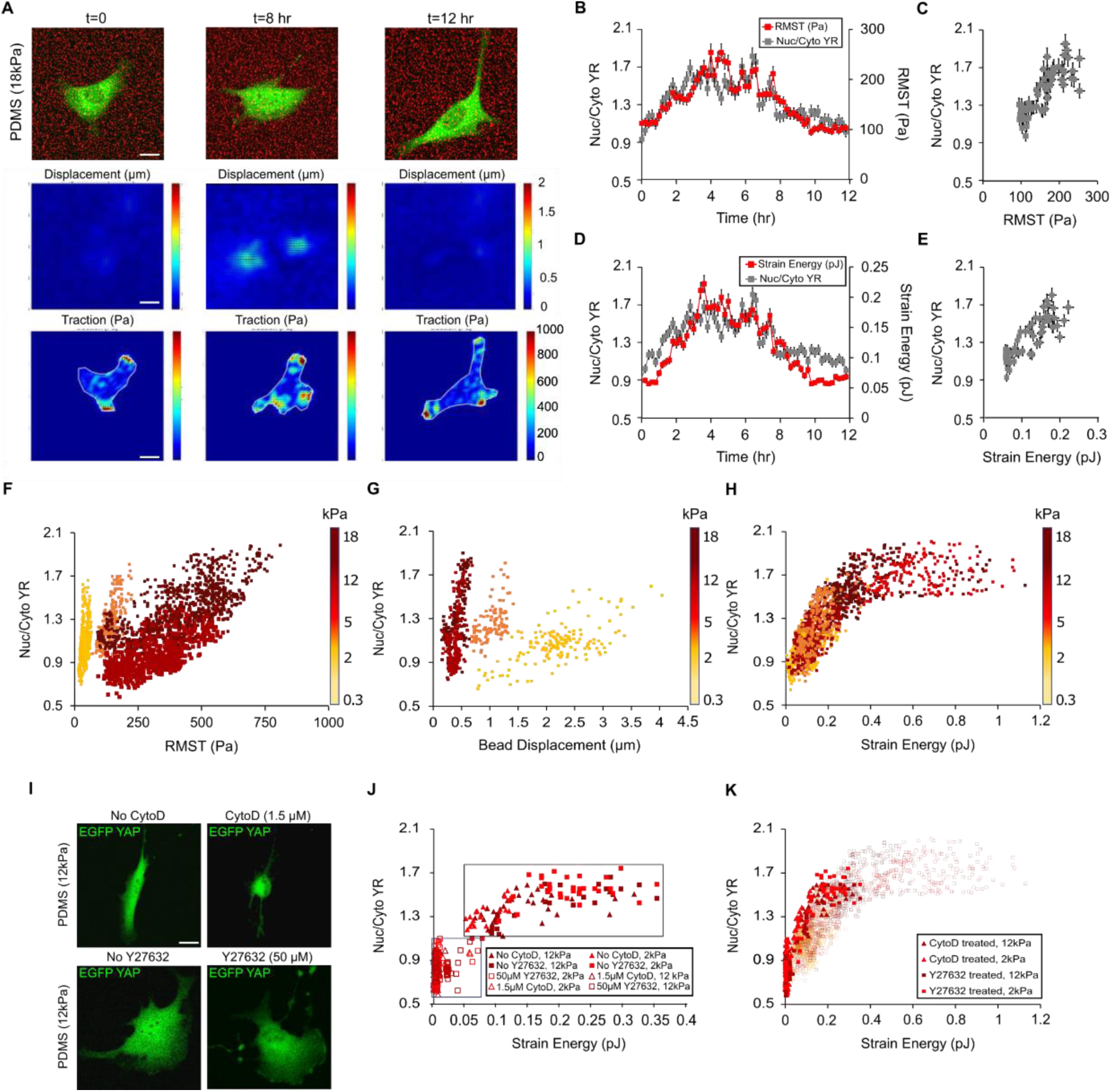
YAP localization is correlated with cell contractility. A) Representative traction stress and bead displacement maps of EGFP-YAP transfected NIH 3T3 cell during its movement on 18 kPa PDMS substrate, B) YR vs RMST over time for the same cell as in (A) with time interval of 12 minutes, C) Scatter plot of YR as a function of RMST, D) Quantification of YR vs Strain Energy for the same cell over time, E) Scatter plot of YR vs Strain Energy for the same cell, F) YR vs RMST for multiple cells seeded on PDMS substrates with different stiffnesses (n>20 cells per condition), G) YR vs bead displacement for the same cells as in (F), H) Scatter plot of YR as a function of Strain Energy for the same cells, I) Example of EGFP-YAP transfected cells before and after pharmacological treatments, J) Quantification of YR vs Strain Energy for multiple cells on PDMS substrates with different Young’s moduli before (solid markers) and 30 minutes after (open markers) pharmacological treatments (n>10 cells per condition), K) All data of YR as a function of Strain Energy for nontreated (partially transparent data) and pharmacologically treated cells. Scale bars are 20 µm. Error bars indicate standard deviation (SD).

We then disrupted myosin activity and the actin cytoskeleton using 50 µM RhoA-associated protein kinase (ROCK) inhibitor (Y27632) and 1.5 µM cytochalasin D (CytoD) (Amano et al., 2010; Mehta and Gunst, 1999) respectively, and monitored their effects on YAP localization (Figures S2I and S2J). Both pharmacological treatments inhibited cell contractility and suppressed YAP nuclear localization ∼15 and ∼40 minutes after CytoD and ROCK inhibitor treatments, respectively (Figures 2I and 2J, Figures S2I and S2J). Critically, these cytoskeletal poisons did not change the relationship between cell contractility and YR, and data from cytoskeleton-disrupted cells followed the same YAP-Strain Energy curve (Figure 2K), suggesting that contractile work predicts YAP localization across all probed cytoskeletal states.

As contractile work appears to determine YAP localization, we considered the cellular components which are mechanically impacted by cell contractile work, and may drive changes in the YR. Since the nucleus is at the heart of YAP activity and has been previously implicated in force-mediated YAP activity (Driscoll et al., 2015; Elosegui-Artola et al., 2017; Kirby and Lammerding, 2018; Shiu et al., 2018), we examined in detail how contractile work mechanically deforms the nucleus and relates to YAP nuclear localization.

### Cell contractility regulates YAP localization via nuclear deformation

To examine the interplay between contractility and nuclear deformation, we quantified the nuclear volume using confocal Z-stacks for cells with different contractility. We found a relationship between contractile work and nuclear volume (Figures 3A and 3B). One key connection between the cytoskeleton and nucleus is the LINC complex (Bouzid et al., 2019; Driscoll et al., 2015; Elosegui-Artola et al., 2017; Lammerding, 2011; Lombardi et al., 2011). To examine the role of the LINC complex in transducing contractile work into nuclear compression, we disrupted the LINC complex by transfecting the cells with two dominant-negative EGFP-Nesprin1-KASH and EGFP-Nesprin2-KASH plasmids (DNK1/2) at the same time iRFP-YAP was transfected (Figure 3A). Lombardi et al showed that overexpression of DNK1/2 inhibits interaction between nesprins and SUN proteins, key components of the LINC complex, at the nuclear envelope by nonspecific binding to endogenous SUN proteins (Lombardi et al., 2011). Our results demonstrated that suppressing the LINC complex in DNK1/2 transfected cells decreased the effect of cell contractility induced nuclear compression (Figure 3B).

**Figure 3.**
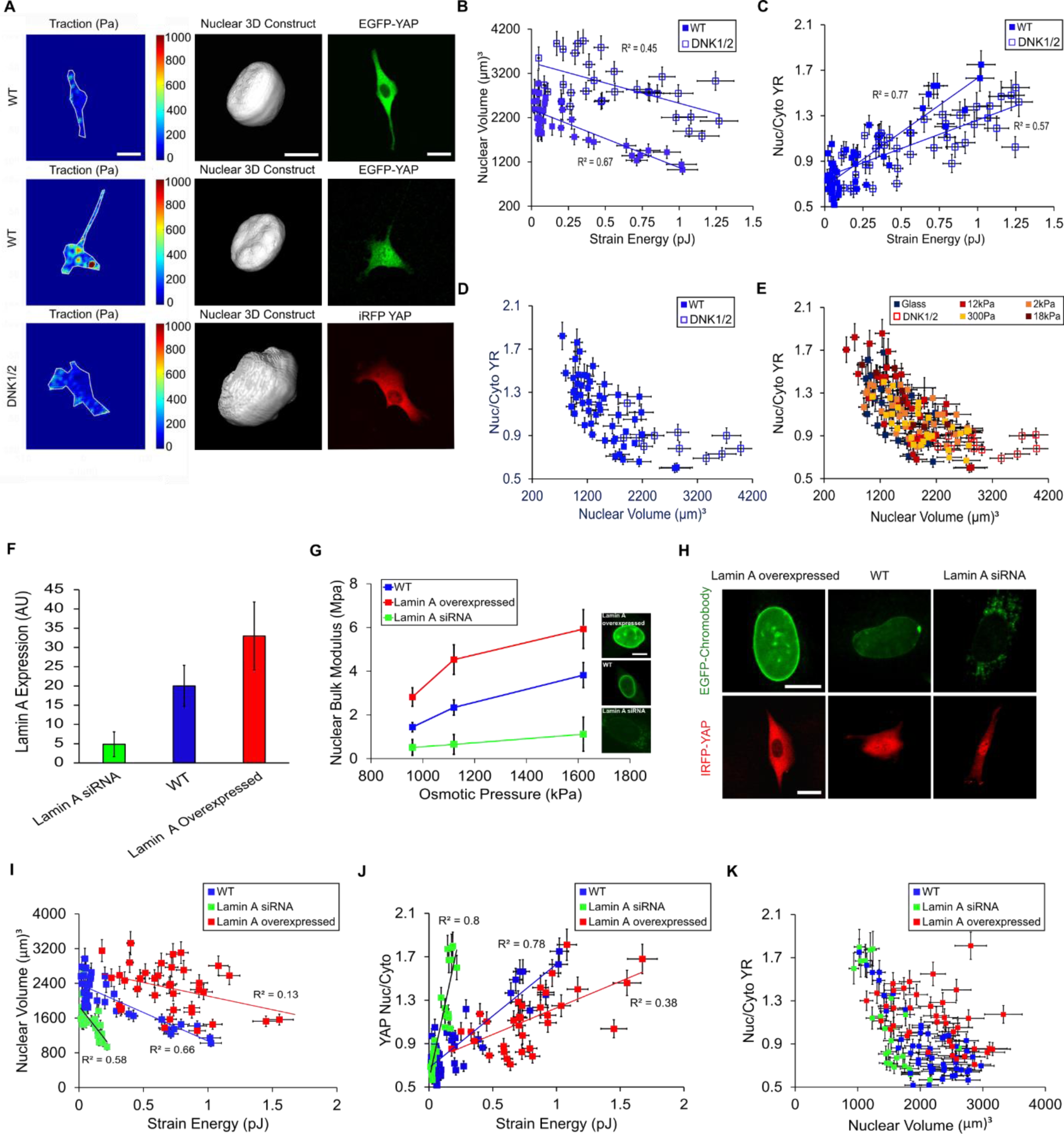
Strain Energy mediated nuclear deformation and YAP activity is directed by LINC complex and nuclear mechanics. A) Representative nuclear volumes and EGFP-YAP distributions in WT and DNK1/2 overexpressed cells with diverse contractility states on 12 kPa PDMS substrate, B) Quantification of nuclear volume vs Strain Energy for WT and DNK1/2 cells seeded on 12 kPa PDMS (n>10 cells per each condition), C) Quantification of YR vs Strain Energy for the same cells as in (B), D) YR as a function of nuclear volume for the same cells, E) Correlation between YR and nuclear volume for WT and DNK1/2 overexpressed cells seeded on PDMS substrates (n>10 cells per each condition), F) Quantification of lamin A expression level for lamin A siRNA, lamin A overexpressed and WT cells (n>20 cells per each condition), G) Quantification of nuclear bulk moduli under different osmotic pressures applied to lamin A siRNA, lamin A overexpressed and WT cells (n>20 cells per each condition), H) Examples of GFP-chromobody and iRFP-YAP in different cells with different lamin A expression level, I) Quantification of nuclear volume vs Strain Energy for cells with different lamin A expression levels (n>15 cells per each condition), J) YR vs Strain Energy for the same cells as in (I), K) YR vs nuclear volume for the same cells. Scale bars are 20 µm for cells and 10 µm for nuclei. Error bars indicate standard deviation (SD).

We then examined how the presence of LINC-mediated contractility influences YAP localization. Here we found that YRs were generally decreased in LINC disrupted cells relative to wild-type (WT) cells for any given cell contractility (Figure 3C, Figures S3A and S3B). This paralleled our observation of the effect of LINC complex disruption on nuclear compression, suggesting a connection between nuclear volume and YR. When we examined YRs as a function of nuclear volume, we found a complete overlap between WT and DNK1/2 transfected cell data, suggesting that nuclear volume may directly regulate YAP activity (Figure 3D). These findings suggest that acto-myosin contractility and LINC-mediated cytoskeletal-nuclear coupling thus contribute to nuclear compression, which appears to describe contractility-based YAP localization.

Returning to the possible influence of substrate stiffness on mechanotransduction mechanisms, we measured YRs as a function of nuclear volume on diverse PDMS substrates with different stiffnesses. Again, we found that cell contractility-driven nuclear compression is correlated with YAP nuclear localization in a substrate stiffness independent way, with similar trends being observed between WT and LINC disrupted cells (Figure 3E). We also observed no trend between nuclear volumes and substrate stiffness (Figure S3C).

Our results highlight the role of nuclear volume and deformation in mechanosensing, which is consistent with the idea that nuclear deformation is necessary to open nuclear pores, allowing YAP nuclear translocation (Elosegui-Artola et al., 2017). We also demonstrated that inhibition of contractile force transfer to the nucleus decreases YAP nuclear import, suggesting that the amount of stress applied to the nucleus moderates its deformation, and consequently YAP activity. We then postulated that varying nuclear stiffness would also modulate contractile-dependent nuclear deformation and YAP localization. This means that the nucleus’ stiffness is likely key in determining specific deformation-mediated mechanosensation to applied stresses.

### Stiffer nuclei require more contractile work to trigger YAP nuclear localization

The interplay between lamin A expression level and nuclear stiffness led us to question whether changes in lamin A-mediated nuclear stiffness would result in changes in contractile work mediated nuclear compression, and corresponding changes in YAP localization. We manipulated lamin A expression levels and measured corresponding nuclei deformation and nuclei stiffness under different osmotic compressions applied by different concentrations of 400 Da polyethylene glycol (PEG) (see Methods) (Guo et al., 2017) (Figures 3F and 3G, Figure S3D). To quantify total single-cell lamin A expression levels, we transfected the cells with GFP-lamin A chromobody (Zolghadr et al., 2012). We found that nuclear stiffness increases as a function of lamin A expression (Figure 3G, Figure S3D).

We then examined how contractile work deforms nuclei with different stiffnesses, and how this in turn regulates YAP localization. We found that in lamin A silenced cells with compliant nuclei (Figure 3G), nuclei were observed to be smaller due to contractility induced nuclear compression (Figure 3I) with increased YRs (Figures 3H and 3J). However, in lamin A overexpressed cells, more contractile work was required to compress the nucleus and localize YAP to the nucleus (Figures 3H-3J, Figure S3E). Since lamin A expression determines the nuclear stiffness, this suggests that contractile work is not a direct regulator of YAP activity, and that nuclear volume may be a more robust independent variable. However, we note that the lamin A overexpressed nuclei deviate from the overall trend of YAP dependence on nuclear volume (Figure 3K).

### YAP translocation correlates with nuclear deformation independent of contractile work, actin filaments and LINC complex

Having observed a correlation between YAP localization and contractility-modulated nuclear volume under the diverse conditions of cytoskeletal poisons, LINC suppression, and varied lamin A expression, we then postulated that modifying nuclear volume through *any* mechanism may be necessary and sufficient for YAP nuclear localization. To test this hypothesis, we applied external osmotic forces on the cells using different concentrations of PEG400, which has previously been shown to reversibly compress cells and nuclei (Guo et al., 2017; Khavari and Ehrlicher, 2019). We then measured the nuclear volumes and YRs for WT cells before and after applying different osmotic pressures, finding the same conserved relationship between nuclear volume and YAP localization (Figures 4A and 4B). Nuclear volume and YRs did not change under 960 kPa pressure, whereas increase in YR and decrease in nuclear volume were observed under 1.62 MPa (Figures 4A and 4B, Figures S4A and S4B). We then considered the role of substrate adhesion in YAP localization by applying osmotic compression to cells in suspension under 10% PEG, which in adhered cells consistently increased YRs.; in compressed suspended cells we did not measure increases in YRs, suggesting that substrate adhesion is an essential aspect of YAP localization in nuclear compression (Figures S4C and S4D).

**Figure 4.**
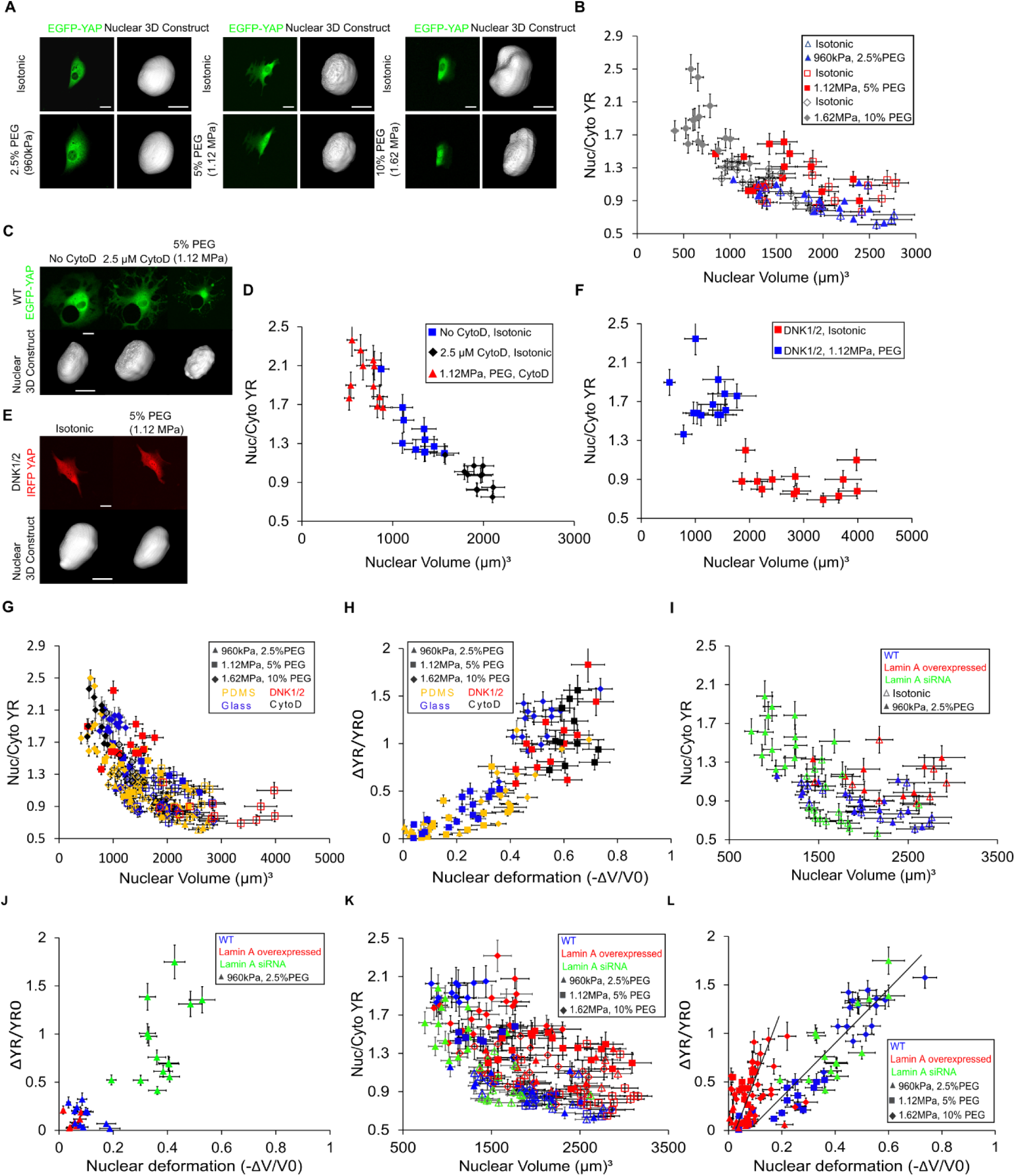
YAP translocation correlates with nuclear deformation independent of contractile work, actin filaments and LINC complex. A) Example of YAP localization and nuclear volume in EGFP-YAP transfected cells under different osmotic pressures, B) Quantification of YR vs nuclear volume in isotonic (open markers) and relevant hyperosmotic conditions (solid markers) (n>10 cells per each condition), C) Example of changes in nuclear volume and EGFP-YAP localization in CytoD treated cells after applying 5% PEG400 (1.12 MPa osmotic pressure), D) YR vs nuclear volume in isotonic condition (blue squares), 30 minutes after adding 2.5 µM CytoD (black diamonds) and 20 minutes after adding 5% PEG400 (red triangle) (n>10 cells), E) Example of changes in iRFP-YAP localization and nuclear volume in GFPDNK1/2 transfected cells before and after adding 5% PEG400, F) Quantification of YR vs nuclear volume for GFP-DNK1/2 transfected cells before (red squares) and after (blue squares) adding 5% PEG400 (n>10 cells), G) All data of YR vs nuclear volumes for CytoD treated (black markers), DNK1/2 (red markers) transfected cells seeded on the glass and WT cells seeded on the glass (blue markers) and 300 Pa PDMS (yellow markers) under different hyperosmotic conditions. All open markers are representative of the cells in isotonic condition and solid markers are the same cells after osmotic compression (n>10 cells per each condition), H) Quantification of YR change 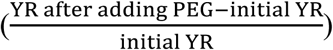 as a function of nuclear volumetric deformation 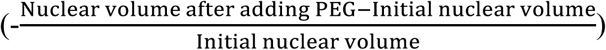 for the same values measured in (G), I) YR vs nuclear volume for WT, lamin A siRNA, and lamin A overexpressed cells before (open markers) and after applying 960 kPa osmotic pressure (solid markers) (n>10 cells per each condition), J) YR change vs nuclear deformation for the same condition as in (I), K) Quantification of YR vs nuclear volume for WT, lamin A siRNA, and lamin A overexpressed cells before (open markers) and after adding applying different osmotic pressures (solid markers) (n>10 cells per each condition), L) Quantification of YR change vs nuclear deformation for the same condition as in (K). Scale bars are 20 µm for the cells and 10 µm for the nuclei. Error bars indicate standard deviation (SD).

Next, we examined the role of the actin cytoskeleton in adhered cells in osmotic pressure mediated YAP activity by depolymerizing F-actin with CytoD. Here we again found the same nuclear volume and YR trend, demonstrating that actin cytoskeleton is not essential for externally compressive YAP mechanosensing (Figures 4C and 4D). Nuclear volumes increased and YRs decreased after depolymerizing actin filaments; however, 1.12 MPa osmotic pressure was sufficient to translocate YAP into the nucleus in the absence of actin (Figure 4D). We also revisited the role of the LINC complex in YAP nuclear localization under external pressure by blocking LINC complex via overexpression of DNK1/2. Compared to WT cells, a lower osmotic pressure (1.12 MPa) was required to significantly deform the nucleus and activate YAP in LINC disrupted cells, demonstrating that external compressive forces deform the nucleus and activate YAP independent of the LINC complex, but that the LINC complex may serve a mechanoprotective role in compression (Figures 4E and 4F). Critically, all YAP activity and nuclear volume data for WT, CytoD treated and DNK1/2 transfected cells appear as a single correlated distribution independent of experimental conditions, demonstrating an apparent connection between nuclear volume and YAP activity (Figure 4G).

While YAP activity appears related to nuclear volume, we questioned how relative nuclear deformation influences changes in YAP activity. To examine this, we measured the change in YAP ratios (ΔYR/YR_0_) as a function of the change of nuclear volume (-ΔV/V_0_) under different external pressures applied to WT, DNK1/2, and CytoD treated cells. For all conditions, the change in YR as a function of change in nuclear volume fell on a single curve, suggesting that nuclear deformation is related to YAP localization in a magnitude-dependent and conserved manner (Figure 4H). We also observed that the nuclei of DNK1/2 and CytoD treated cells deformed relatively more under same amount of pressure as compared with WT cells, further suggesting a potential mechanoprotective role for LINC and the actin cytoskeleton in the context of external compression (Figure 4H, Figure S4E). These results suggest that while acto-myosin contractile work is an essential regulator of YAP under typical cellular conditions, that nuclear deformation underlies YAP localization. This would suggest that nuclear deformability impacts required stress to trigger YAP nuclear transport.

### Lamin A mediated nuclear stiffness regulates the amount of stress required to trigger YAP nuclear transport

We next examined the role of nuclear stiffness modulated by lamin A expression level, finding that cells with increased lamin A expression required more applied stress to reach an equivalent YR. SiRNA lamin A cells required lower pressure (960 kPa) than WT or lamin A overexpressed cells to compress nuclei and localize YAP in the nucleus (Figures 4I and 4J), while 1.12 MPa was required to trigger YAP nuclear localization in WT and lamin A overexpressed cells (Figures 4K and 4L, Figures S4F and S4G) and increased further under 1.62 MPa osmotic pressure (Figures 4K and 4L, Figures S4H and S4I). These findings suggest that lamin A-mediated nuclear stiffness affects the amount of stress required to activate YAP.

These findings describe a preserved relationship between YAP and nuclear volume, observed under different pressures when modifying the stiffness of the nucleus via lamin A expression (Figure 4K). However, we noted that lamin A overexpression itself also appeared to impact YAP localization, with lamin A overexpressed cells yielding higher YAP ratios than those observed in similar volume in WT cells (Figures 4I and 4K, Figures S4F and S4H): lamin A overexpression also led to larger than expected changes in YAP ratios as a function of nuclear deformation (Figure 4L, Figures S4G and S4I). These data suggest that lamin A may play a role beyond that of modulating nuclear stiffness and may be directly related to YAP mechanosensing.

### Lamin A redistributes from the nuclear membrane to nucleoplasm under nuclear deformation and directly regulates YAP localization

When we looked more closely at the data, we found that cells with larger contractility had more compressed nuclei with more YAP in the nucleus as previously established, however, we also noted that the lamin A distribution was impacted; cells with lower contractility had more lamin A in the nuclear membrane (nucleus 1 in Figures 5A, 5C and 5D), whereas cells with larger contractility had a more uniform distribution of lamin A in their nuclei and relatively less in the nuclear membrane (nucleus 2 in Figures 5B-5D). This apparent connection between lamin A localization and contractility prompted further TFM experiments; here we measured the lamin A fluorescence in the nuclear membrane (Lm) as a function of cell contractile work, finding that strongly contractile cells have lower Lm values(Figure 5E), and smaller nuclear volumes (Figure S5A). Additionally, examining YAP localization, we found an inverse relationship between Lm and YR (Figure 5F). We postulated that if contractile forces change the nuclear volume, they may impact lamin A localization, which in turn might directly regulate YAP localization. To test this idea, we quantified Lm as a function of varied nuclear volume by applying different osmotic compressions to WT, lamin A siRNA, and lamin A overexpressed cells. Similar to YAP activation, in lamin A siRNA cells lower force (960 kPa) was sufficient to redistribute lamin A from the nuclear membrane to the nucleoplasm (decrease in Lm) due to high nuclear deformability (Figure 5G). In WT and lamin A overexpressed cells, lamin A stayed intact in the nuclear membrane under the same pressure (Figure 5G), but higher force (1.12 MPa) triggered reduction of lamin A at the nuclear membrane (Lm) of WT and lamin A overexpressed cells (Figure S5C). Minimal Lm values obtained under 1.62 MPa as a result of nuclear deformation (Figure S5G). These results were similar to force mediated YAP activation in different cells with different nucleus’ stiffnesses (Figures 4I and 4K, Figures S4F and S4H).

**Figure 5.**
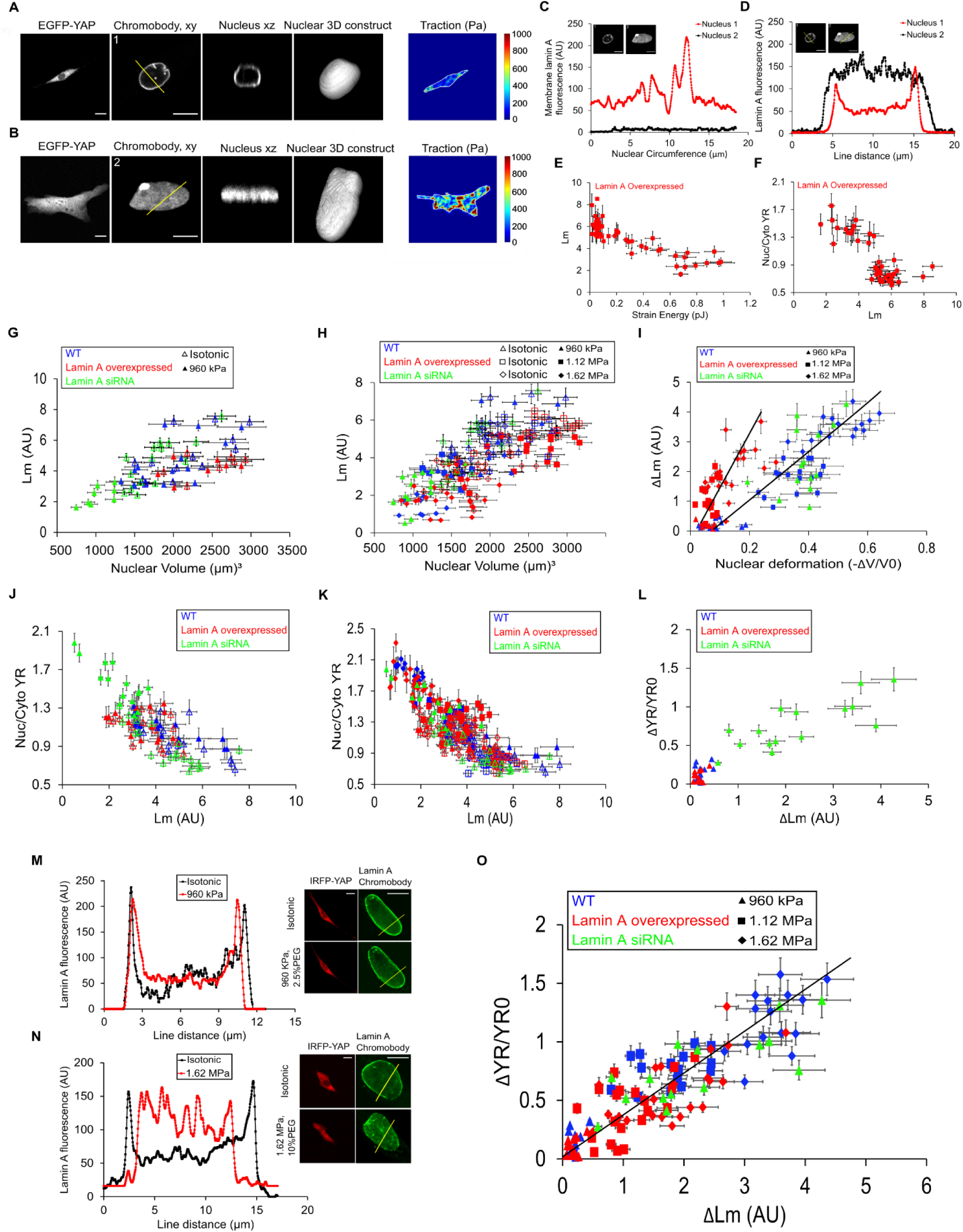
Lamin A redistributes from the nuclear membrane to nucleoplasm under deformation and directly regulates YAP localization. A) Example of YAP localization in low contractile lamin A overexpressed cell with high nuclear volume and high lamin A localized in the nuclear membrane, B) Example of YAP nuclear localization in high contractile lamin A overexpressed cell with more flattened nucleus and even distribution of lamin A throughout the nucleus, C) Quantification of total lamin A fluorescence at the nuclear membrane for nucleus 1 (red line) and nucleus 2 (black line) shown in (A) and (B), D) Lamin A fluorescence along a chord crossing nucleus 1 and 2 in (A) and (B), E) Lm for lamin A overexpressed cells vs Strain Energy (n>15 cells), F) YR vs Lm for the same cells as in (E), G) Lm vs nuclear volume for WT, lamin A siRNA, and lamin A overexpressed cells before (open markers) and after (solid markers) applying 960 kPa pressure (n>10 cells per each condition), H) Collating all data of Lm vs nuclear volume for WT, lamin A siRNA and lamin A overexpressed cells under different experimental conditions (n>15 cells per each condition), I) Collating all data of Lm change (initial Lm − Lm after adding PEG) as a function of nuclear deformation for the same cells as in (H), J) YR vs Lm for WT, lamin A siRNA, and lamin A overexpressed cells before (open markers) and after applying 960 kPa pressure (solid markers) (n>10 cells per each condition), K) Collating all data of YR vs Lm before (open markers) and after (solid markers) adding different concentrations of PEG for the same cells as in (H) (n>10 cells per each condition), L) YR change vs Lm change for WT, lamin A overexpressed and, lamin A siRNA cells under 960 kPa pressure for the same cells as in (G) (n>10 cells), M) Representative of YAP localization and lamin A fluorescence along a chord crossing nucleus of an example cell before (black line) and after (red line) applying 960 kPa pressure, N) YAP localization and lamin A fluorescence along a chord crossing nucleus of another cell before (black line) and after (red line) applying 1.62 MPa pressure, O) Collating all data of YR changes vs Lm changes (ΔLm) for WT, lamin A siRNA, and lamin A overexpressed cells under same conditions as in (K) (n>15 cell per each condition). Scale bars show 10 µm for the nuclei and 20 µm for the cells. Error bars indicate standard deviation (SD).

We found a conserved trend of decreased Lm as a function of decreasing nuclear volume (Figure 5H). However, we noted that the lamin A overexpressed cells deviate from the trend and display lower Lm values for a given volume than WT and lamin A siRNA cells. To further analyze if lamin A does indeed redistribute in response to nuclear deformation, we compared the change in Lm (ΔLm) with the amount of nuclear deformation under osmotic compression. We found a similarly conserved trend under 960 kPa pressure (Figure S5B), whereas under higher pressures we observed a deviation of the lamin A overexpressed cells from WT and lamin siRNA cells (Figure 5I, Figures S5D and S5H) that is reminiscent of our observations in nuclear deformation mediated YAP activation (Figures 3K, 4K, 4L, and 5H). These deviations and similarity between nuclear deformation induced lamin A redistribution and YAP activation cumulatively suggest that YAP is influenced by nuclear volume, but that the lamin A distribution may be a direct independent variable in regulating YAP.

To examine how the lamin A distribution regulates YAP, we plotted YR as a function of Lm, and we found a strong correlation under different osmotic conditions (Figures 5J, Figures S5E and 5SI). When we collated all data, we found a strikingly conserved relationship between lamin A localization and YR, independent of lamin A expression, osmotic pressure, or even nuclear volume (Figure 5K). In lamin A overexpressed and WT cells, under 960 kPa osmotic pressure Lm remains intact with negligible redistribution and YAP remains cytoplasmic (Figure 5M). However, under 1.62 MPa pressure membrane Lm is remarkably redistributed followed by YAP localization (Figure 5N). These findings suggest that the lamin A distribution might be a central regulatory variable for YAP localization.

To further determine if lamin A redistribution is indeed driving YAP translocation, we quantified how changes in Lm due to nuclear compression impacts YAP activity, finding a strong correlation between lamin A redistribution (ΔLm) and YAP activity under diverse osmotic conditions (Figure 5L, Figures S5F and S5J). Collating all data of osmotic compression, we found a pronounced relationship between YR changes and lamin A redistribution for all examined experimental conditions (Figure 5O), suggesting that lamin A redistribution describes YAP nuclear localization independent of nuclear stiffness mediated by overall lamin A expression levels. These findings shed new light on mechanosensing mechanism mediated by nuclear deformation and lamin A reorganization.

## Discussion

Diverse studies have examined how YAP activity is impacted by mechanical stimuli ranging from substrate stiffness to applied forces (Aragona et al., 2013; Dupont et al., 2011; Elosegui-Artola et al., 2017; Nardone et al., 2017; Wang et al., 2016). Here we demonstrated how contractility modulates nuclear deformation and revealed a direct relationship between YAP mechanosensation and lamin A redistribution. Furthermore, we showed that in contrast to previous studies (Aragona et al., 2013; Das et al., 2016; Elosegui-Artola et al., 2017; Fischer et al., 2016), substrate stiffness does not determine YAP activity on PAA or PDMS surfaces. We showed that only in very small rounded cells is YAP principally cytoplasmic (Figure 1G), whereas in spread cells YAP is dynamically distributed between the cytoplasm and the nucleus, independent of substrate stiffness (Figures 1E and 1G). Examining cell contractility, we found that YAP activity appeared correlated with work regardless of substrate stiffness. Furthermore, we showed that contractility varies during cell movement, that YAP activity and contractility are highly correlated temporally, leading to dynamic localization of YAP in single cells independent of substrate moduli (Figures 2B and 2D).

We measured that contractile work compresses the nucleus in a LINC complex mediated way. This nuclear compression in turn regulates YAP localization, which is consistent with the idea of nuclear pore complex opening mediated by nuclear tension (Elosegui-Artola et al., 2017) (Figures 3B-3E). We also varied nuclear stiffness by modulating lamin A expression levels, which changes the amount of nuclear compression under applied physical forces. We found that for a given cell contractility, cells with stiffer nuclei had lower nuclear compression, larger nuclear volume, and lower YRs compared to those with softer nuclei (Figures 3I-3K). These findings suggest that nuclear deformation specifically rather than applied stress mediates YAP localization. Moreover, our results are consistent with previous reports of lamin A overexpression and nuclear stiffening decreasing YAP nuclear transport (Elosegui-Artola et al., 2017; Harada et al., 2014; Swift et al., 2013). However, we observed unexpectedly higher YAP nuclear localization in larger nuclear volumes in lamin A overexpressed cells (Figures 4K and 4L), leading us to examine the interplay between YAP localization and the lamin A distribution.

We found that lamin A localization is mechanosensitive, and that either contractile work or osmotic force induced nuclear deformation causes lamin A to delocalize from the nuclear membrane and to enter the nuclear interior. Recent studies also observed varied localization of A-type lamins with some at the nuclear periphery and some in the nuclear interior depending on cell cycle, differentiation and mechanical cues (Buxboim et al., 2014; Dechat et al., 2010; Gesson et al., 2014; Naetar et al., 2017; Swift et al., 2013; Turgay et al., 2017); however, the pathways involved in nucleoplasmic lamin A regulation and the role of intranuclear lamin A in mechanosening and transcription activation have remained open questions. While the molecular mechanisms behind lamin A disassociation from the nuclear membrane remains unclear, we speculate that this may be related to the local nuclear membrane curvature, and that under high bending curvature that lamin A may delaminate from the nuclear membrane (Figure 6). While nuclear membrane tension has been implicated as a mechanism for YAP regulation, previous work failed to stimulate YAP nuclear localization after applying hypoosmotic solutions that swell nuclei and place the nuclear membrane under tension (Elosegui-Artola et al., 2017); this suggests that nuclear tension alone may not completely describe YAP nuclear localization. Our description of mechanosensitive lamin A redistribution under compression would potentially explain why YAP activation only occurs under nuclear flattening but not under nuclear swelling.

**Figure 6.**
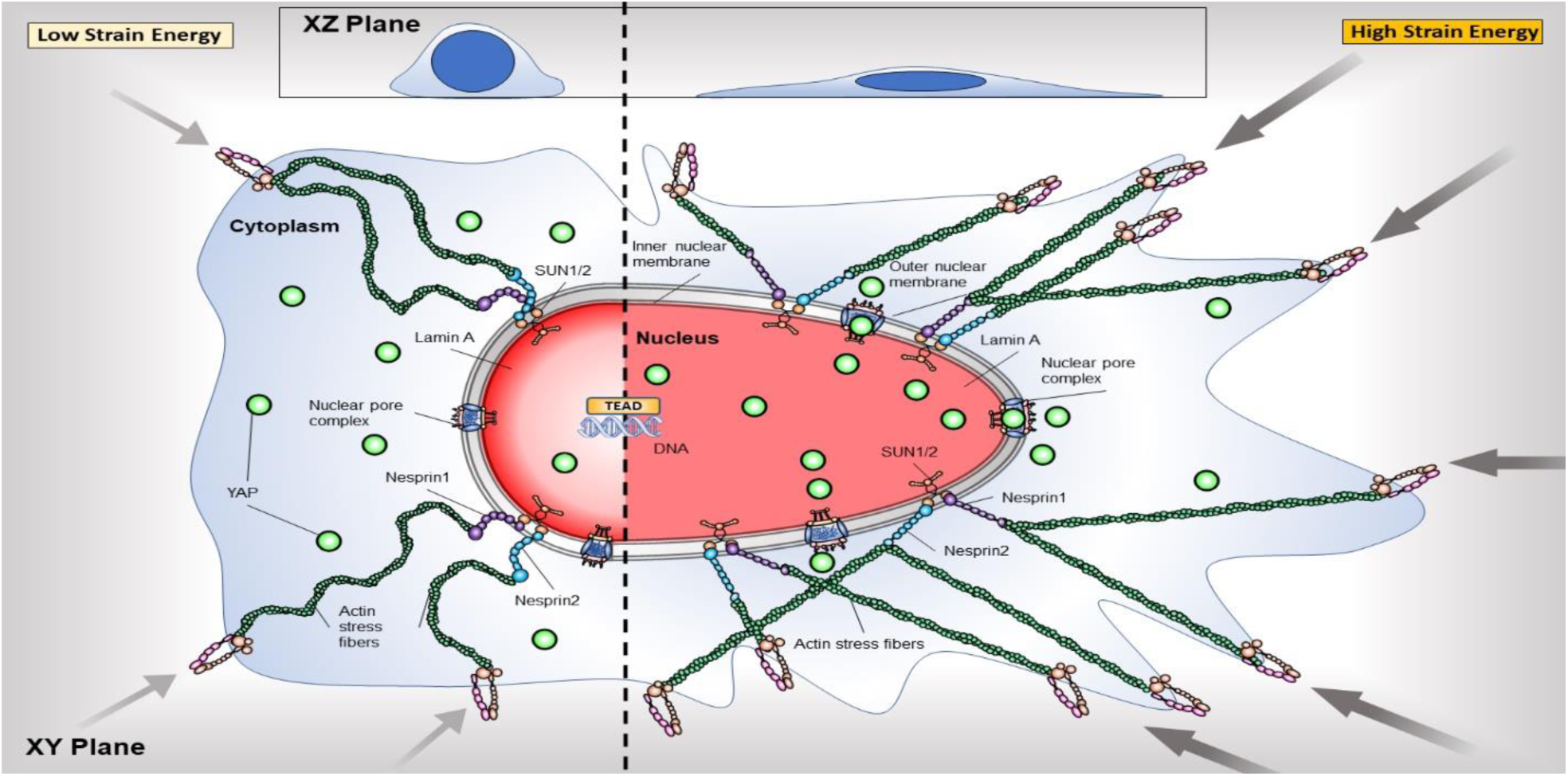
Proposed model regulating YAP nuclear localization. Right part of proposed model is representative of high tensional state of the nucleus with stretched nuclear pores and evenly distributed lamin A (red color) throughout the nucleus, inducing YAP nuclear localization. Left part of the proposed model is representative of lower tension state and less deformed nucleus with lamin A accumulated in the nuclear periphery (darker red) reducing YAP nuclear localization.

Quantifying this redistribution of lamin A from nuclear membrane to nuclear interior, we found that it matched the YAP redistribution in the cell, independent of all other experimental conditions (Figures 5K and 5O). The mechanistic role of lamin A in YAP transport remains unclear, however, it is likely that lamin A interacts with nuclear pore complexes (NPCs). Many studies have reported that lamin A plays a role in regulating the NPC distribution during the cell cycle, and does so in a differentiation dependent way (Guo and Zheng, 2015; Maeshima et al., 2006). Indeed, an inverse correlation between lamin A distribution and NPC density has been observed in the nuclear membrane (Maeshima et al., 2006), suggesting a potential relationship between lamin A localization and YAP translocation, which is consistent with our delamination perspective. Moreover, some transcriptionally active euchromatin regions are associated with nucleoplasmic lamin A (Dechat et al., 2000; Gesson et al., 2016), suggesting that a mechanosensitive redistribution of lamin A may directly impact gene expression and transcription activation.

Our study suggests that nuclear-deformation mediated lamin A reorganization may be a main non-Hippo-dependent YAP regulatory mechanism. This novel mechanism incorporates previously identified mechanical stimuli of YAP regulation, including substrate stiffness, cell contractility, nuclear deformation and nuclear mechanics. This is also consistent with previous reports of lamin A reduction being related to increased YAP activity in cancer progression (Alhudiri et al., 2019; Irianto et al., 2016; Nukuda et al., 2015; Warren et al., 2018; Zanconato et al., 2016a; Zanconato et al., 2016b) We anticipate this link between nuclear stiffness, nuclear deformation, lamin A, and YAP activity will offer new insight and therapeutic strategies for other diverse diseases associated with modified nuclear mechanics and cellular dysfunction, including aging disorders (Capo-Chichi et al., 2011; Verstraeten et al., 2008), Emery-Dreifuss Muscular Dystrophy (Bonne et al., 1999a), and Hutchinson-Gilford progeria syndrome (De Sandre-Giovannoli et al., 2003); A clearer understanding of mechanical YAP regulation may also provide better strategies for directing stem cell engineering and homeostasis. Our proposed interplay between YAP, nuclear volume, and nuclear deformation are consistent with previous reports of cell compression (Bao et al., 2019; Guo et al., 2017; Pan et al., 2018) and high cell contractility (Buxboim et al., 2014; Swift and Discher, 2014; Swift et al., 2013) mediating osteogenic differentiation as a result of YAP activation.

## Acknowledgements

AJE acknowledges support from grants NSERC RGPIN/05843-2014, NSERC EQPEQ/472339-2015, CIHR Grant # 143327, Canadian Foundation for Innovation Projects #32749, #39725. The authors thank Xavier Trepat and Shanahan Catherine for the kind gifts of plasmids (iRFP-YAP, dominant-negative EGFP-Nesprin1-KASH and EGFP-Nesprin2-KASH), and Qiuping Zhang and Johanan Idicula for helpful discussions. The authors sincerely thank Katherine Ehrlicher and Philippe Bergeron for their assistance in copy-editing the manuscript.

## Author contributions

Conceptualization, A.J.E., N.K.; Methodology-development, L.K.S., H.Y.; Methodology-application, N.K.; Plasmid purifications and sequencing, A.G., C.S., L.K.S.; Investigation, N.K.; Writing-original draft, N.K.; Writing-review and editing, N.K., A.J.E., L.K.S., C.S., A.G., C.M.; Funding acquisition-A.J.E.; Resources, A.J.E; Supervision, A.J.E.

## Conflict of interest statement

The authors declare that no conflict of interests exists.

## Materials and Methods

### Fabrication of PDMS substrates

To determine the effects of substrate rigidity on dynamic localization of EGFP-YAP protein, polydimethylsiloxane (PDMS) substrates with different stiffnesses were prepared as described previously (Au - Yoshie et al., 2019; Yoshie et al., 2018). In brief, PDMS solutions were supplied by mixing same weight ratio of component A and B of commercial PDMS (NuSil® 8100, NuSil Silicone Technologies, Carpinteria, CA) with different concentrations of Sylgard 184 PDMS crosslinking agent (Dimethyl, methylhydrogen siloxane, which contains methyl terminated silicon hydride units) to obtain substrates with various stiffnesses (Table 1). Then, 170 μl of each solution was applied to the clean 24*24 mm glass coverslips and cured at 100 °C for two hours. For traction force microscopy, prepared PDMS substrates were coated with 1 µm thick layer of fiduciary particles using spin coater (Laurell Technologies, WS-650 Spin Processor) and incubated at 100 °C for an hour.

**Table 1.** Young’s moduli of different PDMS substrates containing different concentrations of Sylgard 184 crosslink agent.

| Additional crosslinker concentration (weight %) | Young's modulus (kPa) |
| --- | --- |
| 0 | 0.3 ± 0.054 |
| 0.1 | 2 ± 0.062 |
| 0.2 | 5 ± 0.048 |
| 0.36 | 12 ± 0.71 |
| 0.45 | 18 ± 0.68 |
| 1.8 | 100 ± 2.8 |

**Table 2.** Young’s moduli of different PAA hydrogels containing different concentrations of acrylamide, Bis, APS and TEMED.

| (kPa) | Acrylamide % | BIS % | Type | Acrylamide (ul) | BIS (ul) | PBS (ul) | APS (ul) | TEMED (ul) | Beads (ul) |
| --- | --- | --- | --- | --- | --- | --- | --- | --- | --- |
| 1.081 | 3 | 0.1 | AFM | 75 | 50 | 835 | 10 | 10 | 20 |
| 5.008 | 4 | 0.3 | AFM | 100 | 150 | 710 | 10 | 10 | 20 |
| 9.694 | 8 | 0.1 | AFM | 200 | 50 | 710 | 10 | 10 | 20 |
| 15.279 | 8 | 0.15 | AFM | 200 | 75 | 685 | 10 | 10 | 20 |
| 20.85 | 8 | 0.225 | AFM | 200 | 112.5 | 647.5 | 10 | 10 | 20 |
| 23.838 | 8 | 0.3 | AFM | 200 | 150 | 610 | 10 | 10 | 20 |
| 40.397 | 8 | 0.48 | AFM | 200 | 240 | 520 | 10 | 10 | 20 |

### Polyacrylamide Fabrication

Polyacrylamide (PAA) gels were prepared based on previously described protocol (Yeung et al., 2005). PAA gel solutions were prepared with varying concentrations of acrylamide and bisacrylamide mixed with ammonium per sulfate (APS), TEMED and fluorescent fiduciary beads for traction microscopy (Table 1). Acrylamide and bisacrylamide solutions were mixed and degassed for 15-20 minutes under fume hood. Further APS and TEMED were added to the gel solutions and mixed by pipetting. The final solution was added onto the hydrophobic glass slide (treated with RainX) and a coverslip was placed gently on top of the gel drop. After polymerization, the gel sandwiches were placed inside MiliQ water bath and then the glass slides were gently removed.

### Surface modification

In order to covalently bind fibronectin to PDMS or PAA substrates, Sulfo-SANPAH (ThermoFisher Scientific) solution dissolved in 100 mM HEPES was added on top of the substrates and they were exposed to UV for 2 minutes. After UV activation, Sulfo-SANPAH solutions were removed and 5 µg/ml fibronectin (Sigma) solution diluted in PBS was added on to of the samples, followed by incubation in room temperature for 9-12 hours. Finally, fibronectin solutions were removed, and substrates were rinsed with PBS 3 times. After UV sterilization of coated substrates, trypsinized cells were seeded on top of the samples and they were allowed to adhere overnight.

### Cell culture

NIH-3T3 *Mus musculus*, mouse cell line was obtained from ATCC and cultured in Dulbecco’s modified Eagle medium (DMEM) (Wisent) supplemented with 10% fetal bovine serum (FBS) (Wisent) and 1% Penicillin-Streptomycin antibiotic (P/S) (Thermo Fisher). The cells were incubated at 37 °C in 5% CO_2_ environment and, they were allowed to grow on the substrates for 18 hours before imaging.

### Transfection and Confocal microscopy of live cells

To quantitatively track EGFP-YAP mechanotransduction with time, we transiently transfected NIH-3T3 cells with 2 plasmids, pEGFP-yap-C3-hYAP1 (Addgene, plasmid #17843) (Basu et al., 2003) and EBFP2-Nucleus-7 (nuclear localization signal, Addgene, plasmid #55249), using GenJet transfection reagent (Signagen). 18-24 hours later, cells were seed on fibronectin-coated PDMS, PAA and glass substrates and after cell attachment, they were transferred to a lab-built heated stage perfused with 5% CO_2_ and mounted on a confocal microscope (Leica TCS SP8 with a 10x objective). With this setup, we could image cells with transmission and fluorescence microscopy for extended periods, while maintaining a controlled culture environment.

In order to examine the effects of LINC complex on contractile and force mediated nuclear deformation and YAP translocation, we transfected the cells with two dominant-negative GFP-Nesprin1-KASH and GFP-Nesprin2-KASH (DNK1/2) plasmids which were kindly provided by Dr. Catherin Shanahan’s laboratory (King’s College, London) (Lombardi et al., 2011). Previously it has been shown that overexpression of dominant-negative Nesprin-KASH disrupts interaction between nesprins and SUN proteins at nuclear envelop by nonspecific binding to endogenous SUN proteins resulting in mislocalization of nuclear nesprins and disruption of LINC complex (Elosegui-Artola et al., 2017; Lombardi et al., 2011). To specify YAP localization, at the same time we transfected the cells with iRFP-YAP which was a gift from Xavier Trepat (Institute for bioengineering of Catalonia (IBEC), Barcelona) and EBFP-Nucleus to avoid any crosstalk between GFP-DNK1/2 and YAP and 18 hours after transfection we seeded the cells on PDMS substrates coated with fluorescent beads followed by incubation at 37 °C for 12 hours. Finally, alive transfected cells seeded on PDMS traction substrates were imaged overtime using confocal Leica SP8 with 10x objective. To measure traction stress and strain energy at the same time, fluorescent beads coating PDMS and PAA substrates also were imaged along with EGFP-YAP transfected cells and EBFP-Nucleus.

To quantify YAP nuclear to cytoplasmic ratio, image segmentation was performed using matlab code for every time frame to measure the ratio of EGFP-YAP fluorescence inside the nucleus to EGFP-YAP fluorescence in the cytoplasm during cell movement on PDMS, PAA and glass substrates.

### Immunostaining

To compare endogenous YAP localization with EGFP-YAP, we fixed the cells with 4% paraformaldehyde for 15 min in room temperature and washed 3 times with PBS. We stained the nuclei with 0.5 µl/ml bisBenzimide H 33342 trihydrochloride (Sigma) and after 20 minutes, cells were washed with PBS. The cells were permeabilized with 0.1% Triton X-100 diluted in PBS for 10 minutes. To avoid any nonspecific hydrophobic binding, 2% bovine serum albumin (BSA) was added to the cells and incubated for 30 minutes in room temperature. After washing with PBS, we made solution of 10 µg/ml of YAP mouse monoclonal antibody (sc101199, Santa Cruz) and donkey Anti-Mouse IgG H&L (Alexa Fluor® 647) secondary antibody (ab150107, abcam) in BSC separately. The cells were separately incubated first with primary antibody and then with the secondary antibody for an hour in room temperature. Fluorescence images were acquired with a Leica SP8 confocal microscope and endogenous YAP ratio values were measured using image segmentation and quantifying the ratio of YAP intensity inside the nucleus to YAP intensity in cytoplasm, and then the values were compared with YR values obtained from EGFP-YAP transfected cell.

### Cell spread area

To examine the effects of cell spread area, we started confocal imaging 10 minutes after seeding the EGFP-YAP and EBFP-Nucleus transfected cells on fibronectin-coated PDMS substrates and continued imaging for 18 hours using confocal Leica SP8 with low magnification (x10 objective). Then we quantified YAP ratio and cell projected area for every time frame acquired by confocal microscope. The projected cell area of the cells was determined with Fiji software.

### Traction Force Microscopy

Active contractile stress in actin cytoskeleton were quantified using Traction Force Microscopy (TFM) as previously described (Yoshie et al., 2018). In brief, EGFP-YAP transfected NIH 3T3 cells were cultured on fibronectin coated compliant PDMS substrates of known moduli and a thin PDMS layer of embedded fiduciary fluorescent particles was spin coated on the top surface. After 12 hours incubation at 37 °C, EGFP-YAP transfected cells and fluorescent particles were imaged simultaneously over time using Leica TCS SP8 confocal microscope with low magnification (x10/NA 0.4 air objective) at a resolution of 0.28 µm/pixel. The reference images of the particles were acquired at the end of the experiment by detaching the cells from the substrate surface.

Cell-substrate traction stresses and strain energies were calculated for each acquired time frame as described previously (Butler et al., 2002). Briefly, local displacements of the fiduciary particles were calculated by comparing the particle positions with cells on the substrate and reference particle positions without cells. From the particle displacement and known Young’s moduli of the PDMS substrate, cellular contractile stresses and strain energies were calculated. Quantifying YAP ratio based of above EGFP-YAP and EBFP-Nucleus images for the relevant time frames, we could measure YAP localization for every contractile state of the cells on PDMS substrates with different stiffnesses.

### Pharmacologic treatments

To further quantify the effects of contractile stress and work (strain energy) on YAP subcellular translocation, we started imaging of the EGFP-YAP transfected cells and fluorescent particles on top of PDMS substrates with different stiffnesses before any treatment and then we added 1.5 µM Cytochalasin D (CytoD, ThermoFisher Scientific) to the cells which depolymerize actin filaments and decrease contractility. We then continued imaging for another 2 hours after adding CytoD followed by killing the cells at the end of the experiment for the reference images required for traction stress analysis. Quantifying YAP ratio and relevant traction stress as well as strain energy for each time frame before and after actin filaments depolymerization enable us to track changes in YAP localization in real time during losing cells’ contractilities in real time. Moreover, we applied 50 µM ROCK inhibitor (Y27632, abcam) to nontreated EGFP-YAP transfected cells seeded on different PDMS substrates to investigate how inhibition of actomyosin activity affects dynamic movements of YAP in alive single cells. We continued imaging for 6 hours after ROCK inhibitor treatment and at the end of experiment the cells were detached for traction analysis.

### Quantification and modulating of lamin A expression

To quantify the total lamin A present in the nuclei, all the cells were transfected with GFP tagged lamin A Chromobody (Chromtek) which facilitates real time imaging of lamin A by labeling the total lamin A without interfering with its function and distribution in the nuclei (Srivastava et al., 2019; Zolghadr et al., 2012). As previously it has been verified that lamin A Chromobody is the accurate quantitative metric of lamin A expression and distribution (Srivastava et al., 2019), we transfected NIH 3T3 cells with GFP-Chromobody to quantify lamin A expression and distribution. To modulate lamin A expression in GFP-Chromobody transfected NIH 3T3 cells, we overexpressed lamin A by transiently transfecting the intact cells with m-Cherry tagged plasmid DNA for lamin A which was a gift from Michael Davison (Addgene, plasmid # 55068). To surpass lamin A expression, we transfected the cells with RFP tagged inducible shRNA construct for lamin A (Dharmacon) by adding 0.5 µg/ml doxycycline to WT cells transfected with GFP tagged lamin A chromobody (Matsushita et al., 2013), followed by incubation at 37 °C for 72 hours.

After modulating lamin A expression, cells were seeded on the fibronectin-coated glasses and after 24 hours incubation they were imaged using confocal Leica SP8 microscope with x63/1.4 NA oil immersion objective. Lamin A expression level in different cells was assessed by identifying a mask covering the whole nucleus and then quantifying total GFP-Chromobody’s fluorescence in the nuclear mask using MATLAB code.

In order to examine lamin A intensity localized in the nuclear membrane, we identified a mask that covered only the nuclear membrane, and GFP-Chromobody’s fluorescence was only quantified in the defined nuclear membrane mask (Lm).

To study the effects of lamin A expression and distribution on YAP localization, all the cells were transfected with iRFP YAP as well as EBFP-Nucleus at the same time when we were transfecting GFP-Chromoboy and modulating lamin A expression. 18 hours after transfection and 72 hours after doxycycline treatment cells were seeded on fibronectin-coated glass and PDMS traction substrates, followed by 24 hours incubation at 37 °C and 5% CO_2_ environment. To quantify YAP ratio, total lamin A expression, lamin A localized in the nuclear membrane and contractility, iRFP-YAP, EBFP-Nucleus, GFP-Chromobody and fluorescent beads were imaged by confocal microscope with 63X/1.4 NA oil immersion objective.

### 3-D volume measurement of nuclei

In order to measure nuclear volume and nuclear deformation, XYZ stacks of EBFP tagged nuclei with a z-step size of 0.5 µm were imaged using Leica SP8 confocal microscope with x63/1.4NA oil immersion objective lens. The 3D visualization and quantification were performed by Fiji software. To quantify nuclear volume, the stacks were theresholded based on the top and bottom of the nuclei and then the number of voxels of the theresholded region was counted and multiplied by the size of each voxel using Fiji software. To investigate the role of cell contractility in nuclear deformation and YAP localization in WT, lamin A overexpressed and lamin A siRNA cells, we employed z stacks imaging of the EBFP tagged nuclei of the single cells seeded on PDMS substrates coated with fluorescent particles at the same time when iRFP-YAP, EBFP-Nucleus, GFP-Chromobody and fluorescent particles underneath of each cell were imaged. To analyze traction stresses and strain energies of the cells, at the end of the experiment, cells were detached for the null force image.

We repeated the same experiment for LINC disrupted cells and CytoD treated cells to investigate how LINC complex disruption and actin filament depolymerization affect contractile force mediated nuclear deformation.

### Osmotic compression

To examine how externally deform nuclei regulated lamin A distribution and YAP activation, hyperosmotic pressure was applied using different concentrations of 400Da polyethylene glycol (PEG 400, Sigma) (Guo et al., 2017; Khavari and Ehrlicher, 2019; Srivastava et al., 2019). First, XYZ stacks of EBFP-Nucleus, iRFP-YAP and GFP-Chromobody of the cells seeded on the fibronectin-coated glasses and PDMS substrates were imaged using confocal microscope with x63/1.4 NA. Then, different concentrations of PEG400 were added to the cells and again z stacks of the nuclei as well as iRFP-YAP and GFP-Chromobody were imaged (Table 3). We then quantified nuclear volume, YAP ratio (YR) and nuclear membrane lamin A (Lm) after adding different concentrations of PEG400 to the cells and compared them with initial values obtained in isotonic condition to investigate how applied force deforms nucleus, redistribute lamin A and activates YAP.

**Table 3.** Quantified osmotic pressures relevant to different concentrations of PEG400.

| PEG concentration wt% | Osmotic pressure (MPa) |
| --- | --- |
| 2.5 | 0.96 |
| 5 | 1.12 |
| 10 | 1.62 |

### Quantification of nuclear bulk moduli

To determine how nuclear deformability influenced the required force to activate YAP, we modulated lamin A expression level as mentioned above and 18 hours after seeding the cells on fibronectin-coated glasses, we applied different hyperosmotic pressures using different concentrations of PEG400. We acquired XYZ stacks of EBFP-Nucleus before and after adding PEG. We then quantified change in the nuclear volume when the cells with different lamin A expression were exposed to different hyperosmotic shocks and using the following equation,

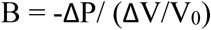

where B= bulk modulus, ΔP = osmotic pressure, ΔV= change in nuclear volume and V_0_ = Initial nuclear volume, we calculated nuclear bulk modulus (Khavari and Ehrlicher, 2019; Srivastava et al., 2019).

