## Supplementary material for "Lamin A redistribution mediated by nuclear deformation determines dynamic localization of YAP": Supplemenat file

**Supplementary materials:**

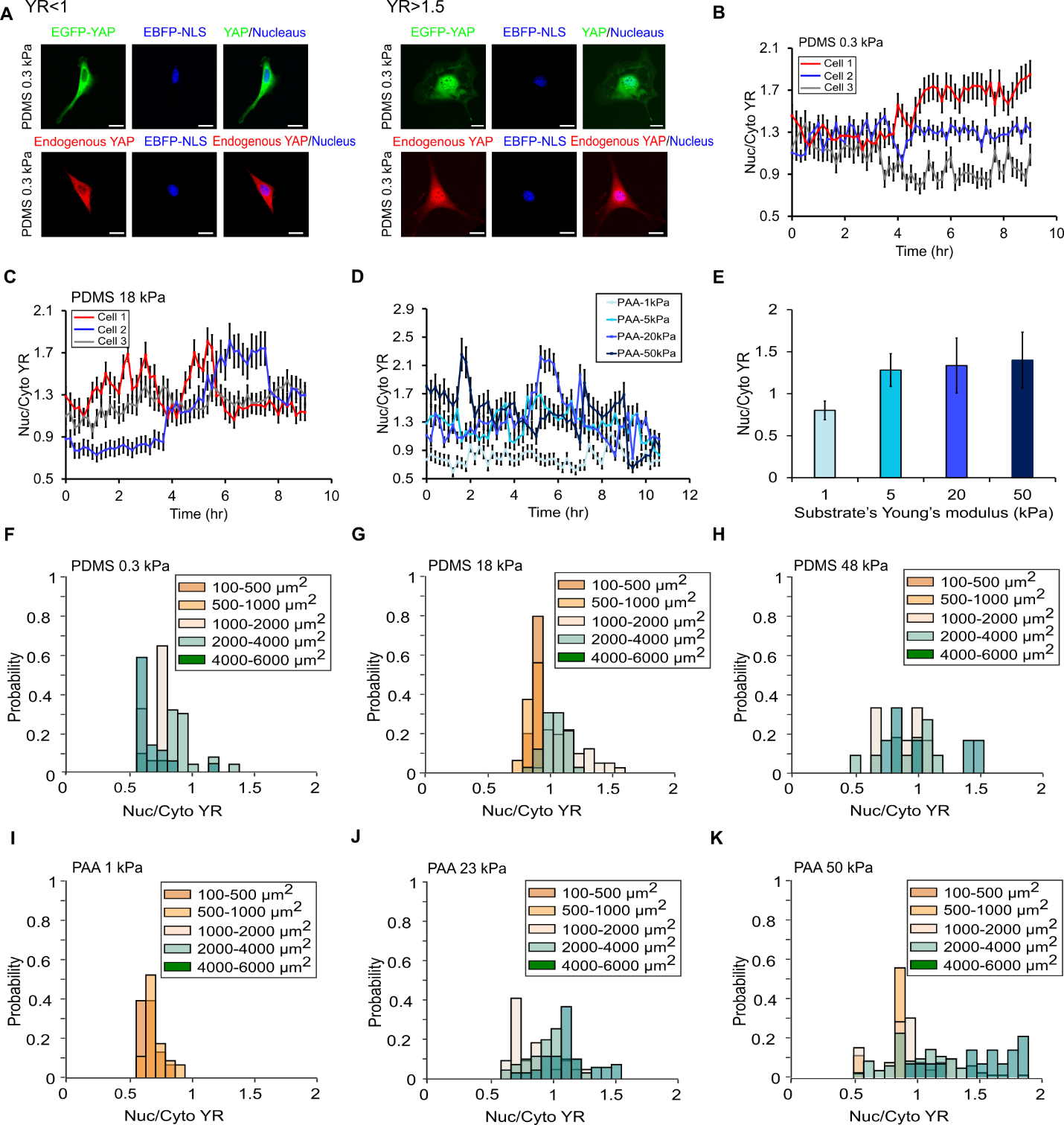

**Figure S1. YAP localization is dynamic overtime and independent of substrate stiffness and cell spread area.** A) Example image of different EGFP-YAP and endogenous YAP localization in NIH 3T3 cells labeled with EBFP-NLS nucleus on soft PDMS with Young’s modulus of 0.3 kPa, B) Quantification of YAP localization variation overtime in 3 different EGP-YAP transfected cells moving on soft PDMS with Young’s modulus of 0.3 kPa, and C) Stiff PDMS with Young’s modulus of 18 kPa, D) Quantification of YAP ratio variation in different EGFP-YAP transfected cells during cell movement on PAA substrates with various stiffnesses, E) Time average of YAP ratio on different PAA substrates for the same conditions as in (D), F) YAP ratio distribution based on cell spread area for different cells seeded on PDMS substrates with Young’s modulus of 0.3 kPa, G) 18 kPa, and H) 48 kPa. Different colors are representative of different range of cell spread area (n>10 per each condition), I) YAP ratio distribution based on cell spread area for different cells seeded on PAA substrates with Young’s modulus of 1 kPa, J) 23 kPa and K) 50 kPa. Different colors are representative of different range of cell spreading area (n>15 per each condition). Scale bars are 20 µm. Error bars indicate standard deviation (SD).

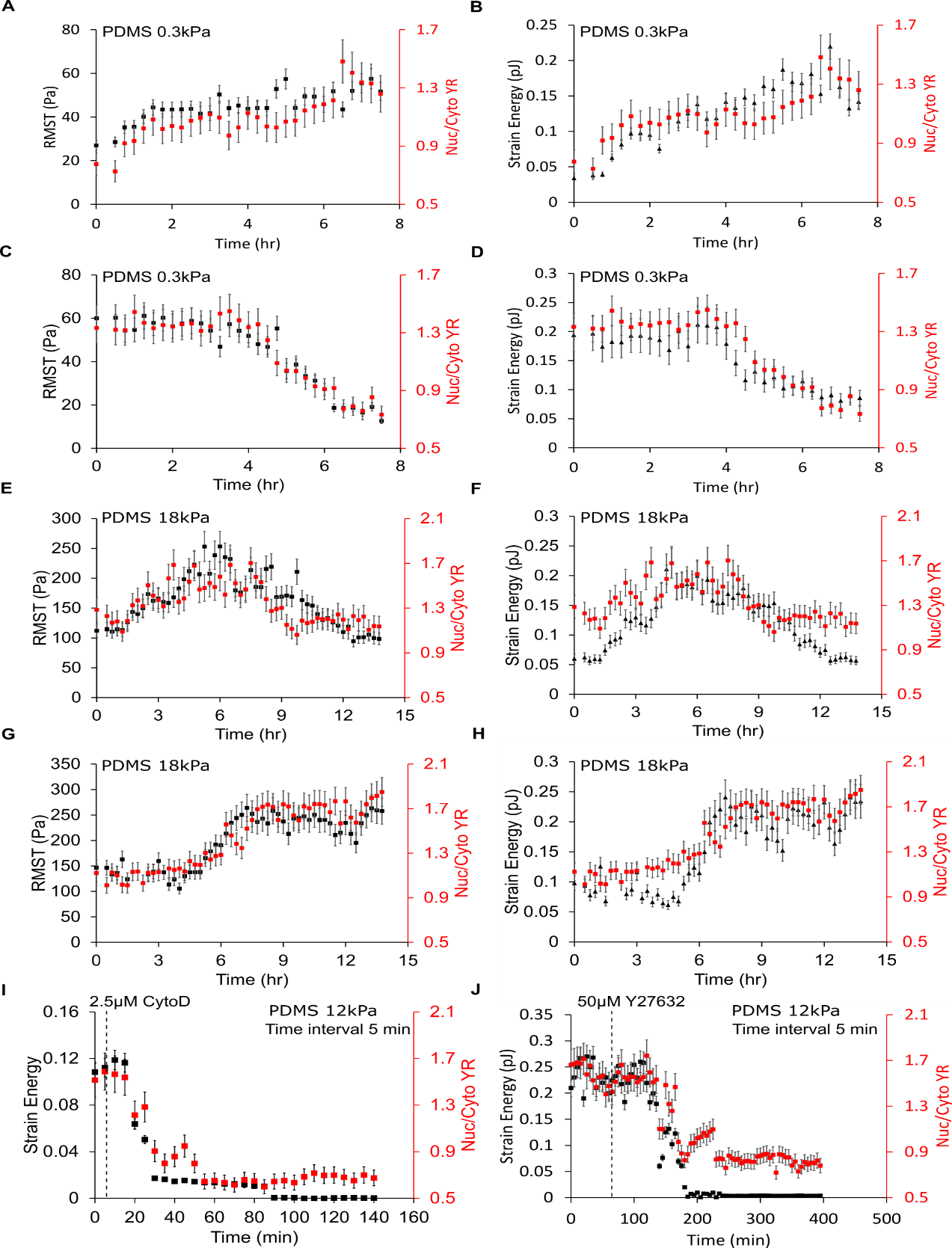

**Figure S2. YR is correlated with cell contractility, best with Strain energy.** A) Example of quantified RMST and EGFP-YAP ratio during cell movement on PDMS substrate with Young’s modulus of 0.3 kPa, B) Quantification of Strain Energy and YAP ratio for the same cell in the same condition as in (A), C) Quantification of RMST and YR for a different cell seeded on PDMS with the same stiffness, D) Strain Energy and YR measured for the same cell in the same condition as in (C), E) Example of tracking RMST and EGFP-YAP localization at the same time during cell movement on PDMS with stiffness of 18 kPa, F) Quantification of Strain Energy and YR for the same cell in the same condition as in (E), G) Another example of tight correlation between YAP ratio and RMST for a different cell seeded on PDMS with stiffness of 18 kPa, H) Strain Energy and YR measured for the same cell in the same condition as in (G), I) An example of YAP cytoplasmic localization as a result of losing contractility due to actin filament depolymerization about 15 minutes after CytoD treatment of EGFP-YAP transfected cells seeded on PDMS with Young’s modulus of 12 kPa, J) An example of decrease in YAP ratio due to actomyosin inhibition about 50 minutes after adding ROCK inhibitor to the EGFP-YAP transfected cells seeded on PDMS with the same stiffness. Error bars indicate standard deviation (SD).

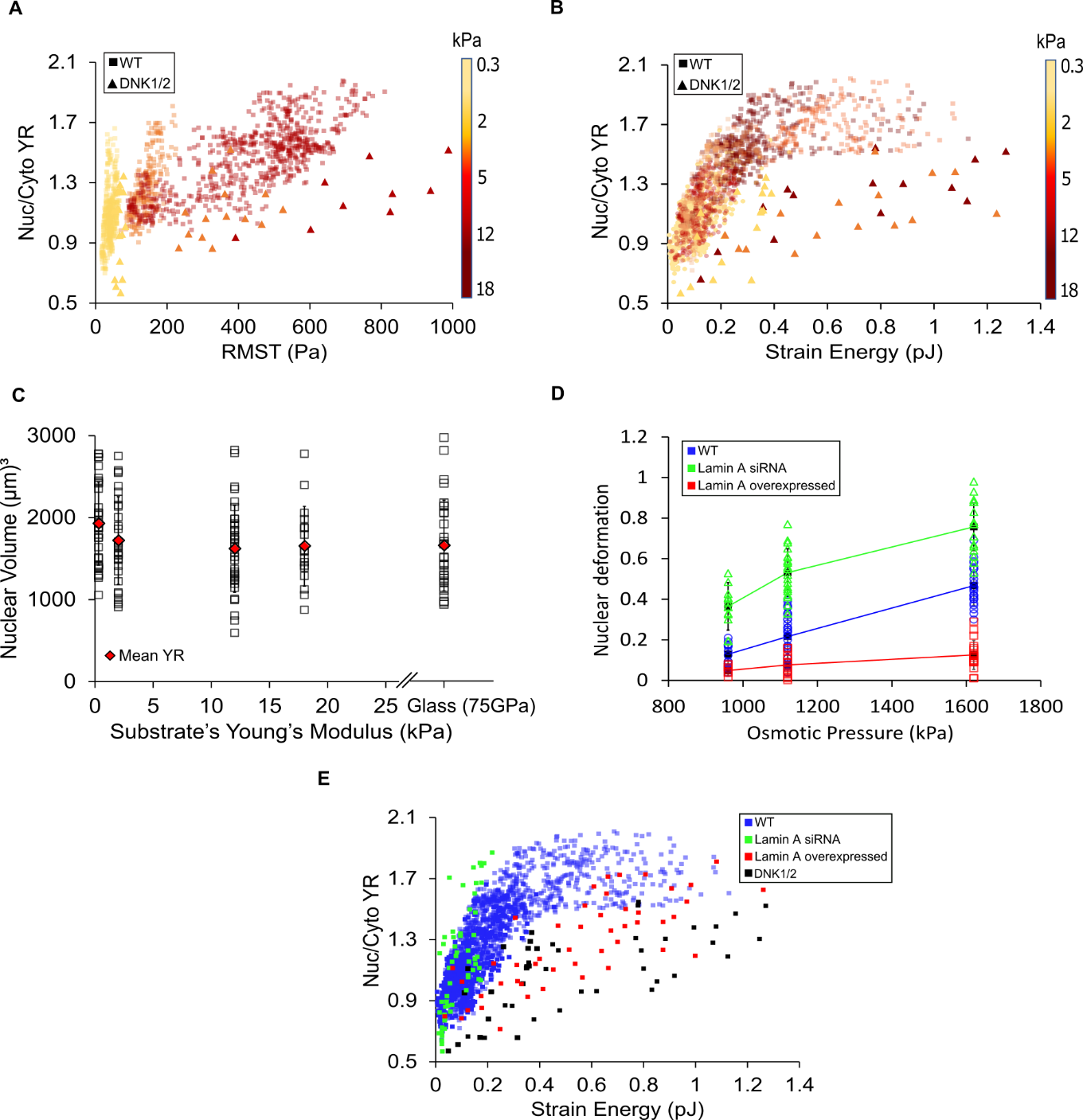

**Figure S3. Strain Energy mediated nuclear deformation and YAP activity is directed by LINC complex and nuclear mechanics.** A) Comparing RMST and YAP ratio for WT (squares) and DNK1/2 (triangles) cells seeded on PDMS substrates with different stiffnesses. Color bar is representative of substrate stiffness (n>15 per each condition), B) Quantification of YAP ratio and Strain Energy for the same cells in the same conditions as in (A), C) Quantification of nuclear volume of different cells seeded on divers PDMS substrates and glass (n>15 per each condition). Diamonds are representative of mean nuclear volume of the cells on each stiffness, D) Quantification of nuclear volumetric deformation (-ΔV/V0) under different osmotic pressures applied by PEG400 to the WT, siRNA lamin A and lamin A overexpressed cells, E) All the data of Strain Energy as a function of YR for WT (blue markers), lamin A siRNA (green markers), lamin A overexpressed (red markers) and DNK1/2 cells (black markers) (n>15 per each condition). Error bars indicate standard deviation (SD).

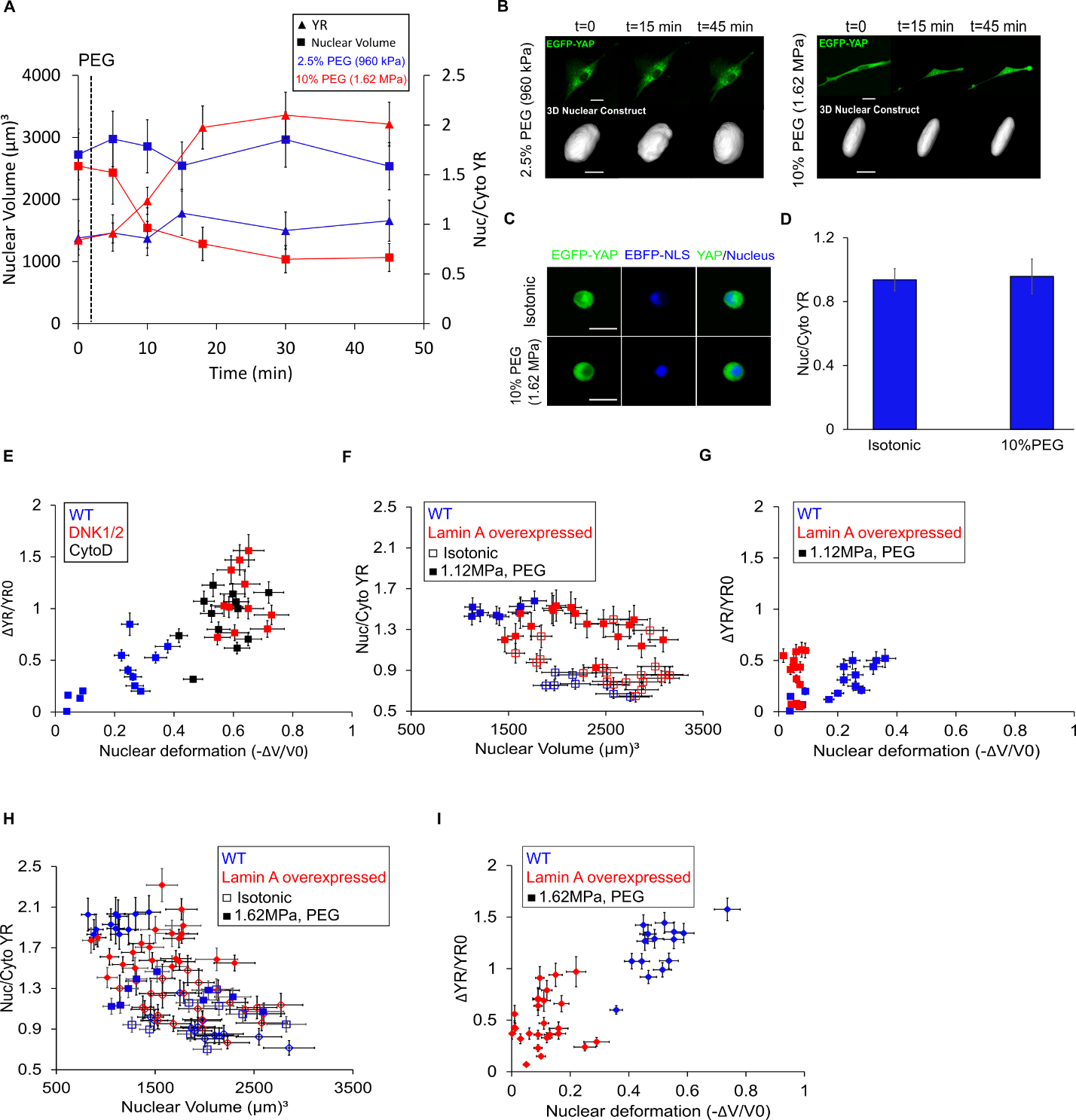

**Figure S4. Nuclear deformability induced by lamin A expression level directly regulates required force to activate YAP.** A) Example of tracking nuclear volume and EGFP-YAP localization overtime in 2 different cells before and after adding 2.5% (blue markers) and 10% PEG (red markers), B) Example image of changes in nuclear volume and EGFP-YAP localization over time under different hyperosmotic conditions (2.5% and 10% PEG). Scale bars show 20 µm for cells and 10 µm for nuclei, C) Example image of changes in YAP nuclear localization in suspended cells transfected with EGFP-YAP. Scale bars show 20 µm, D) Quantification of mean YR values of different suspended cells before and after adding 10% PEG (n>15 cells per each condition), E) Quantification of differential YAP ratio as a function of nuclear deformation for WT (blue markers), DNK1/2 (red markers), and CytoD treated (black markers) cells under 5% PEG (1.12 MPa osmotic pressure) (n>10 cells per each condition, F) Quantification of YAP ratio as a function of nuclear volume for WT (blue markers) and lamin A overexpressed (red markers) cells before (open markers) and after (solid markers) adding 5% PEG (1.12 MPa osmotic pressure) (n>10 cell per each condition), G) Measuring differential YAP ratio as a function of nuclear deformation for the same cells in the same conditions as in (F), H) Quantification of YR as a function of nuclear volume for WT (blue markers) and lamin A overexpressed (red markers) cells before (open markers) and after (solid markers) adding 10% PEG (1.62 MPa osmotic pressure) (n>10 cells per each condition), I) Quantification of differential YAP ratio as a function nuclear volumetric deformation for the same cells in the same conditions as in (H). Error bars indicate standard deviation (SD).

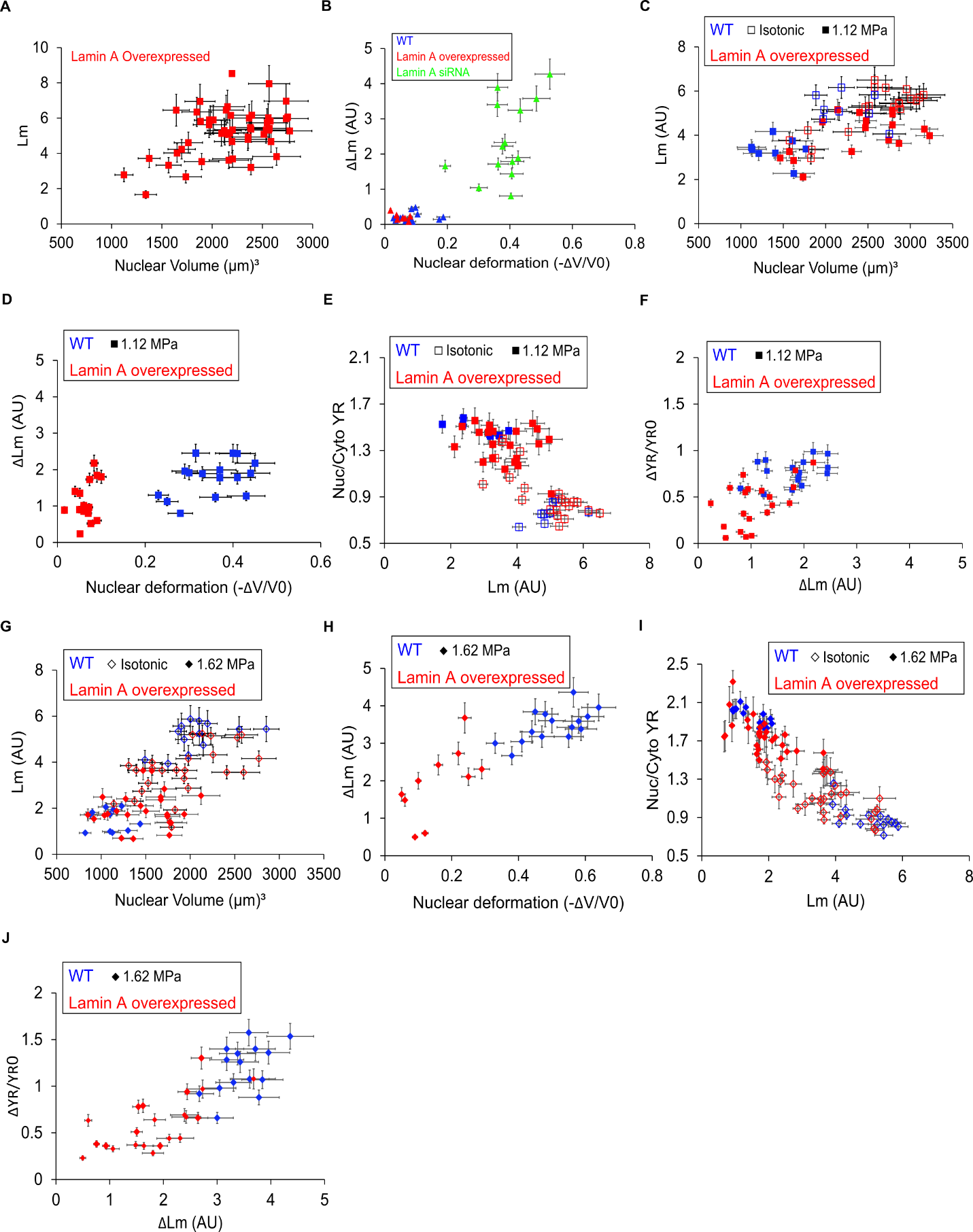

**Figure S5. Lamin A redistributes from the nuclear membrane to nucleoplasm under deformation and directly regulates YAP localization.** A) Quantification of nuclear membrane lamin A as a function of nuclear volume for lamin A overexpressed cells (n>15 cells), B) Quantification of lamin A redistribution ($Initial Lm before PEG-Lm after adding PEG)$ as a function of nuclear deformation for WT (blue), lamin A siRNA (green) and lamin A overexpressed (red) cells under 2.5% PEG (n>10 cells per each condition), C) Quantification of membrane lamin A vs nuclear volume for WT (blue markers) and lamin A overexpressed (red markers) cells before (open markers) and after applying 5% PEG (1.12 MPa osmotic pressure, solid markers) (n>10 per each condition), D) Quantification of change in membrane lamin A as a function of nuclear deformation for the same cells under the same condition as in (C), E) Quantification of YR as a function of Lm for the same cells before (open markers) and after (solid markers) adding 5% PEG, F) Quantification of change in YAP ratio as a function of change in membrane lamin A distribution for the same cells under the same pressure as in (C), G) Assessment of membrane lamin A vs nuclear volume for WT (blue markers) and lamin A overexpressed (red markers) cells before (open markers) and after applying 10% PEG (1.62 MPa osmotic pressure, solid markers) (n>15 per each condition), H) Quantification of lamin A redistribution as a function of nuclear deformation for the same cells under the same condition as in (G), I) Quantification of YR as a function of Lm for the same cells before (open markers) and after (solid markers) adding 10% PEG, J) Quantification of change in YAP ratio as a function of change in lamin A distribution for the same cells under the same pressure. Error bars indicate standard deviation (SD).
